## Supplemental Figures for "AmpliconReconstructor: Integrated analysis of NGS and optical mapping resolves the complex structures of focal amplifications in cancer"

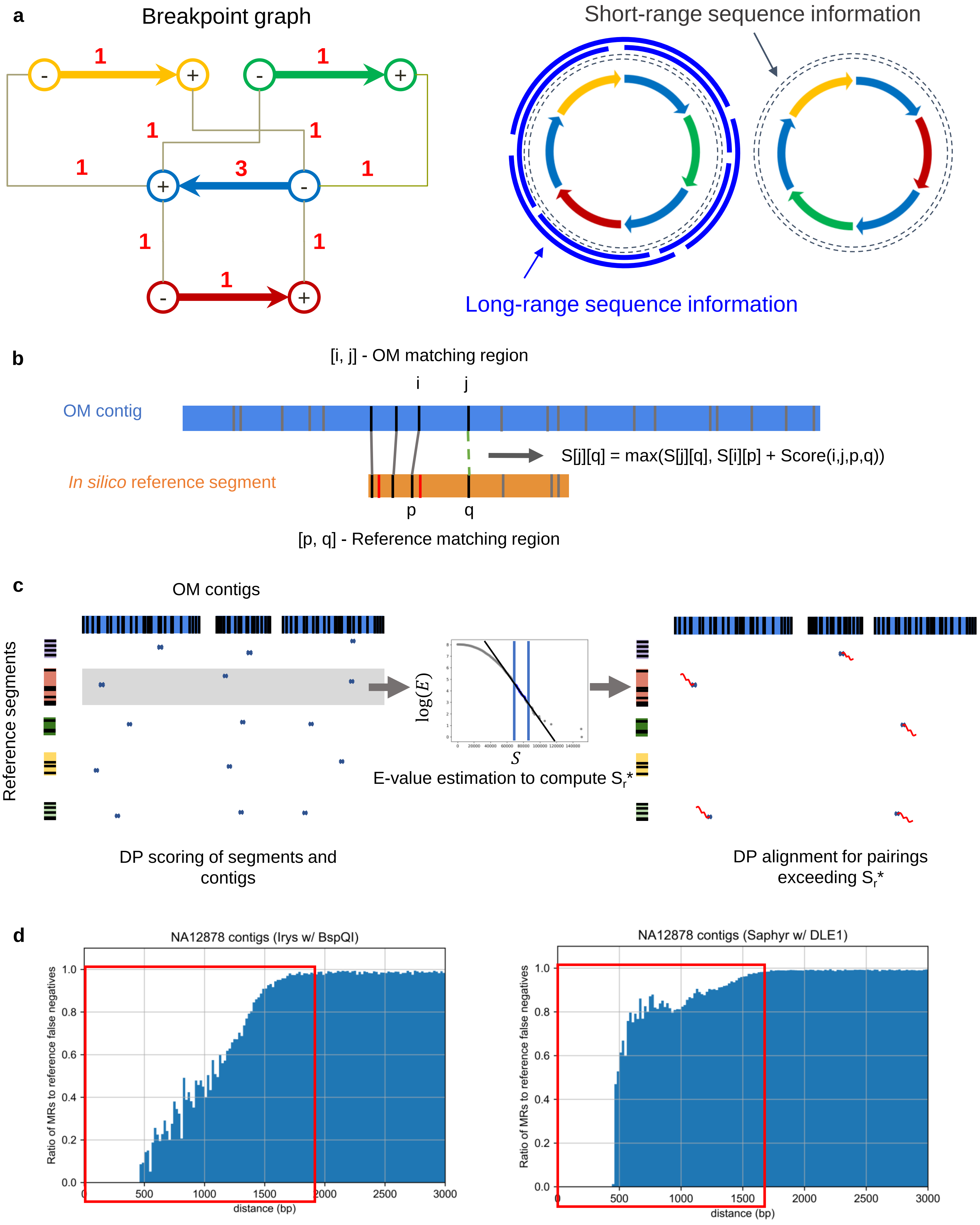
a
Breakpoint graph
Short-range sequence information
Long-range sequence information
b
[i, j] - OM matching region
OM contig
S[j][q] = max(S[j][q], S[i][p] + Score(i,j,p,q))
In silico reference segment
[p, q] - Reference matching region
c
OM contigs
Reference segments
E-value estimation to compute S_r*
DP scoring of segments and contigs
DP alignment for pairings exceeding S_r*
d
NA12878 contigs (Irys w/ BspQI)
NA12878 contigs (Saphyr w/ DLE1)
Ratio of MRs to reference false negatives
distance (bp)

**Figure S1.**

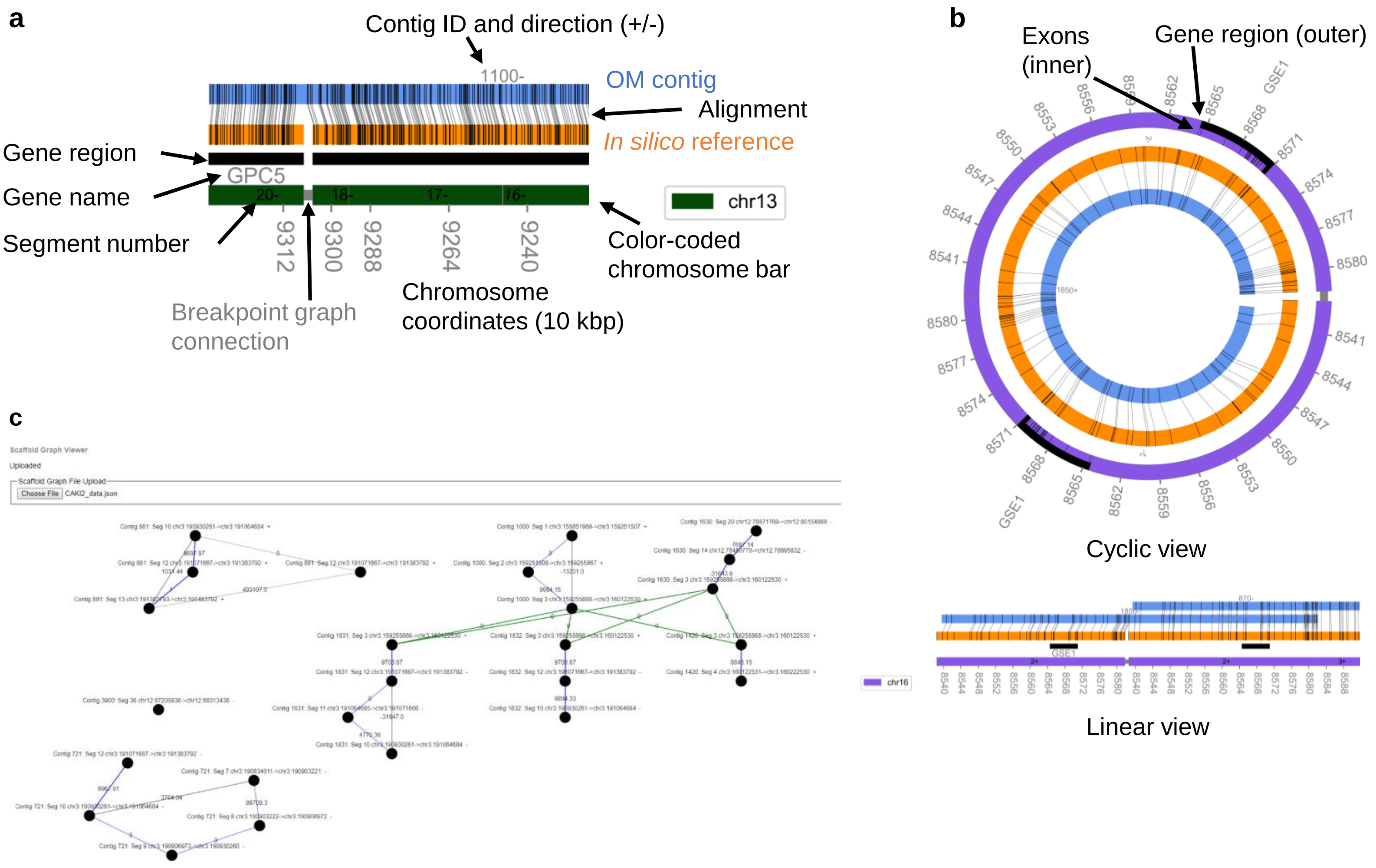
a
Contig ID and direction (+/-)
1100-
OM contig
Alignment
In silico reference
Gene region
GPC5
Gene name
Segment number
20-
18-
17-
16-
chr13
9312
9300
9288
9264
9240
Color-coded chromosome bar
Breakpoint graph connection
Chromosome coordinates (10 kbp)
b
Exons (inner)
Gene region (outer)
GSE1
Cyclic view
Linear view
chr16
c
Scaffold Graph Viewer
Uploaded
Scaffold Graph File Upload
Choose File
CAKI2_data.json

**Figure S2.**

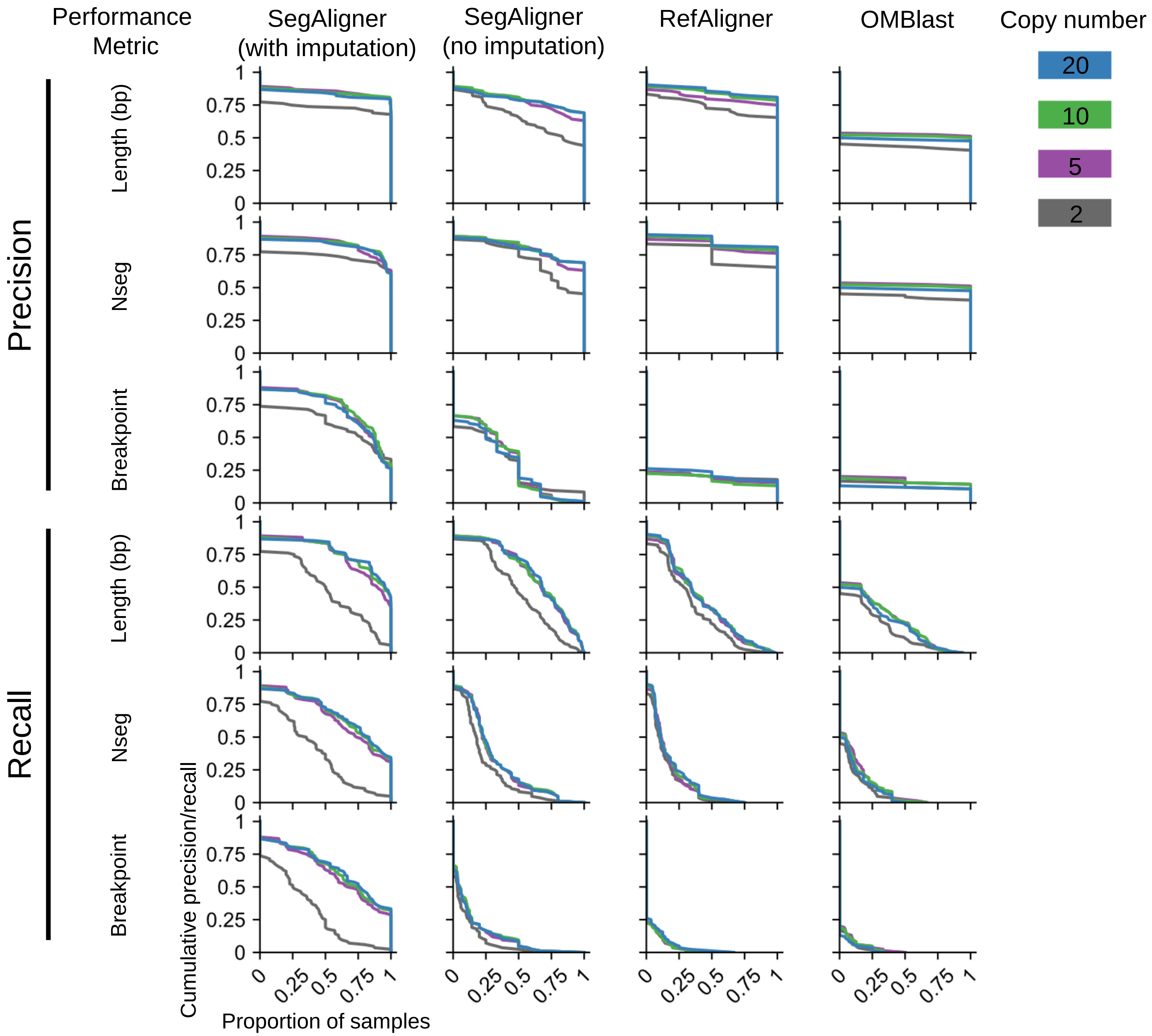
Performance Metric
SegAligner (with imputation)
SegAligner (no imputation)
RefAligner
OMBlast
Copy number
20
10
5
2
Precision
Recall
Length (bp)
Nseg
Breakpoint
Length (bp)
Nseg
Breakpoint
1
0.75
0.5
0.25
0
0.25
0.5
0.75
1
Cumulative precision/recall
Proportion of samples

**Figure S3.**

**a** False edges added

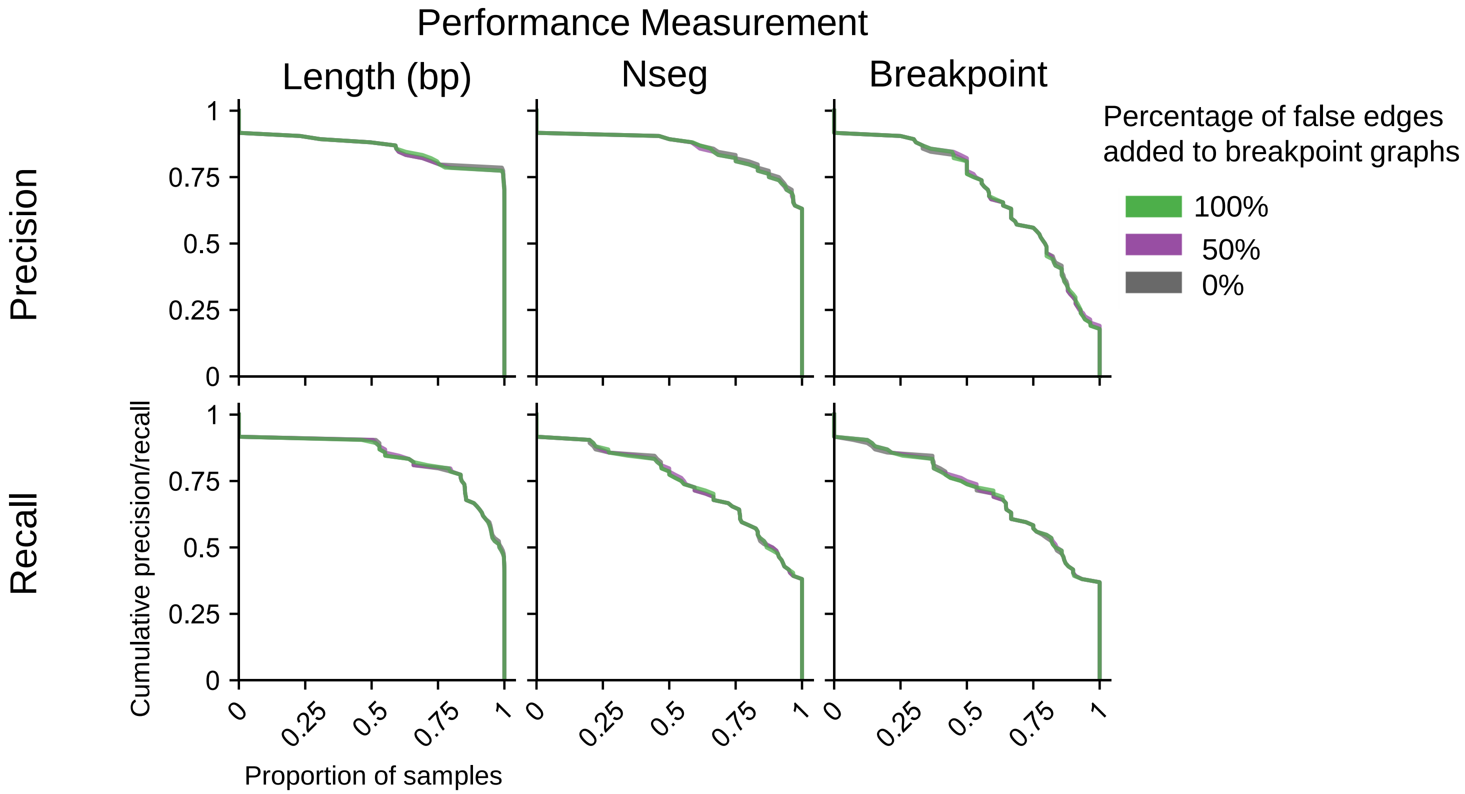


**b** Combination of simulated amplicons

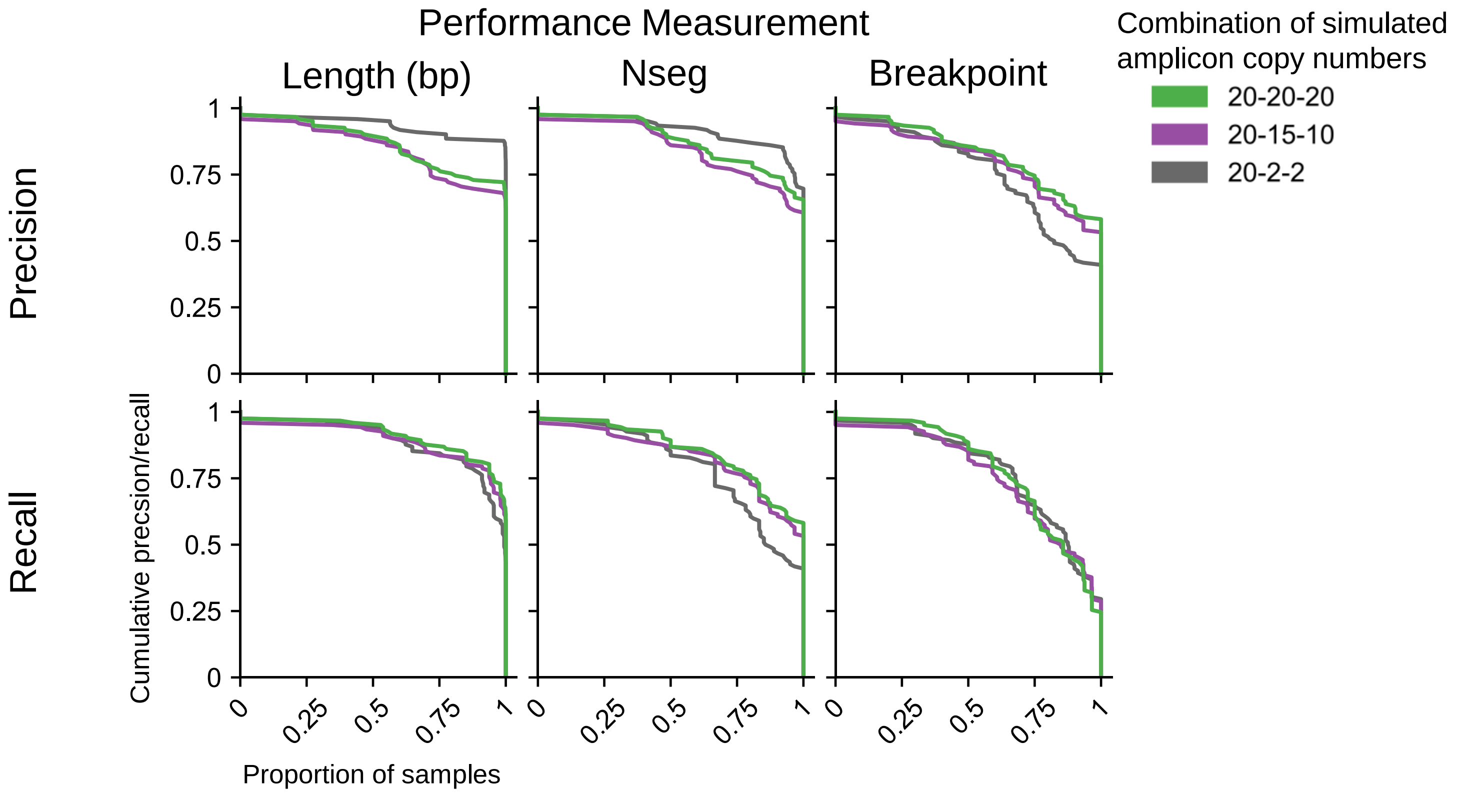


**Figure S4.**

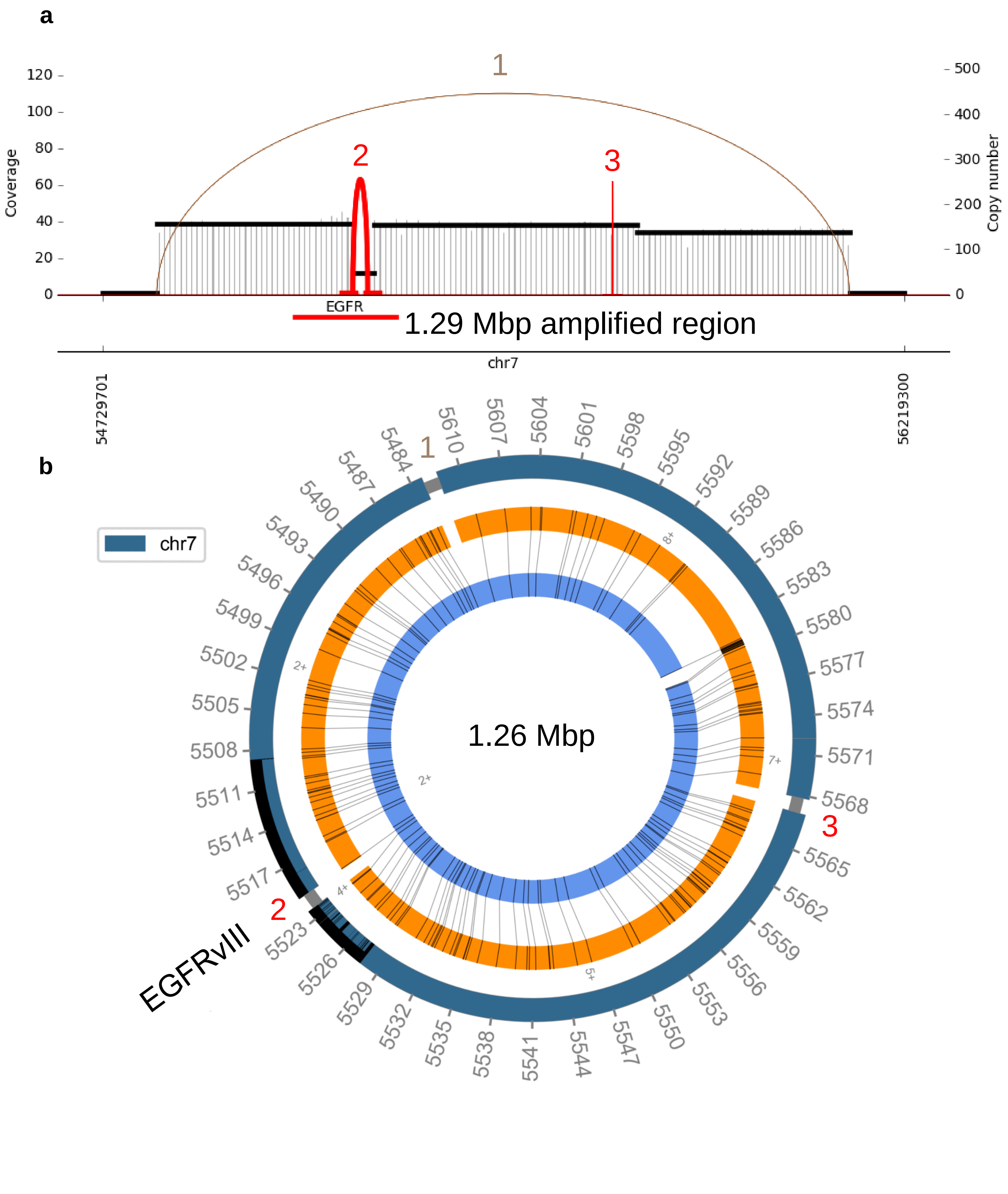
a
b
Coverage
Copy number
EGFR
1.29 Mbp amplified region
chr7
54729701
56219300
chr7
1.26 Mbp
EGFRvIII
1
2
3

**Figure S5.**

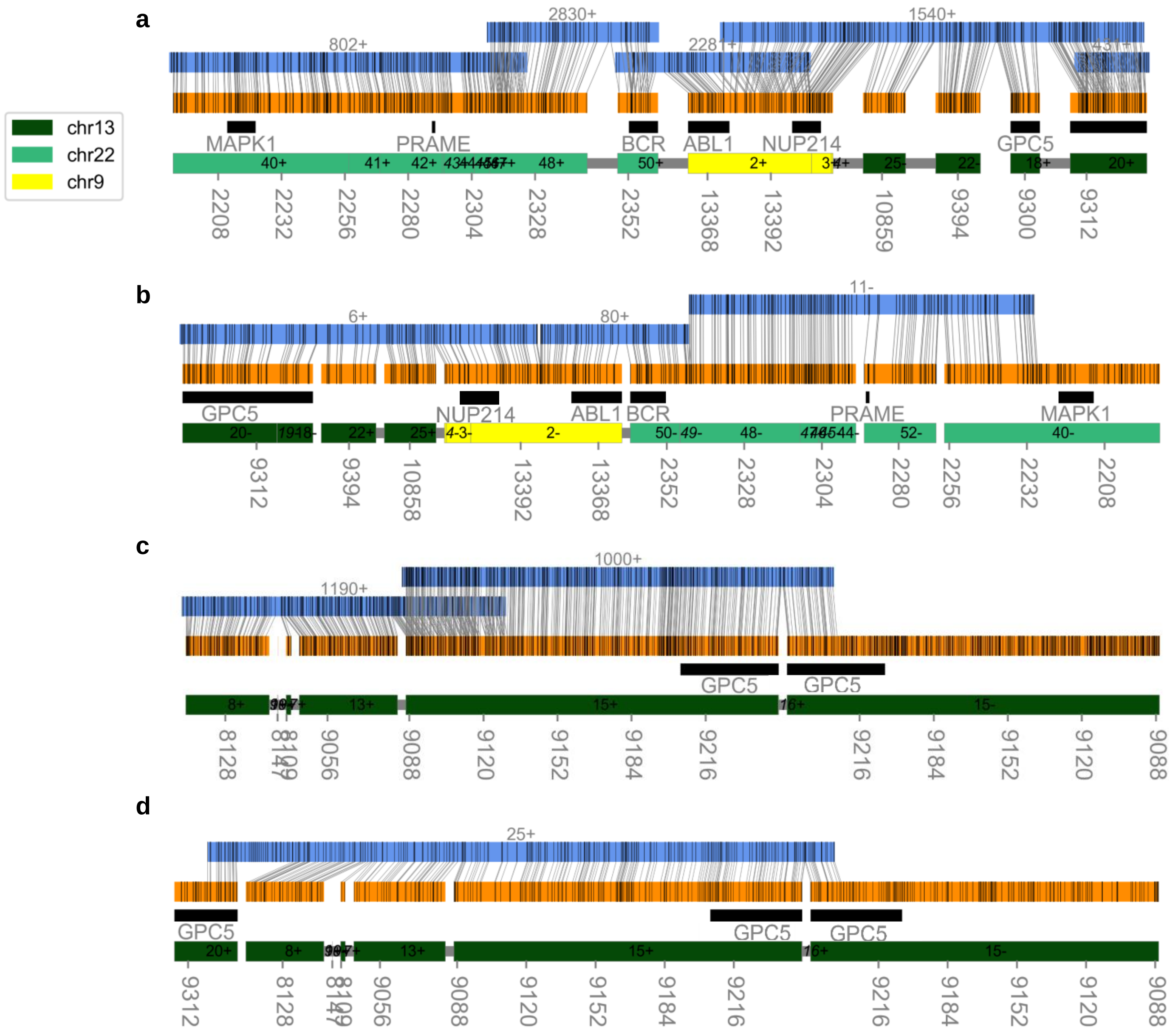


**Figure S6.**

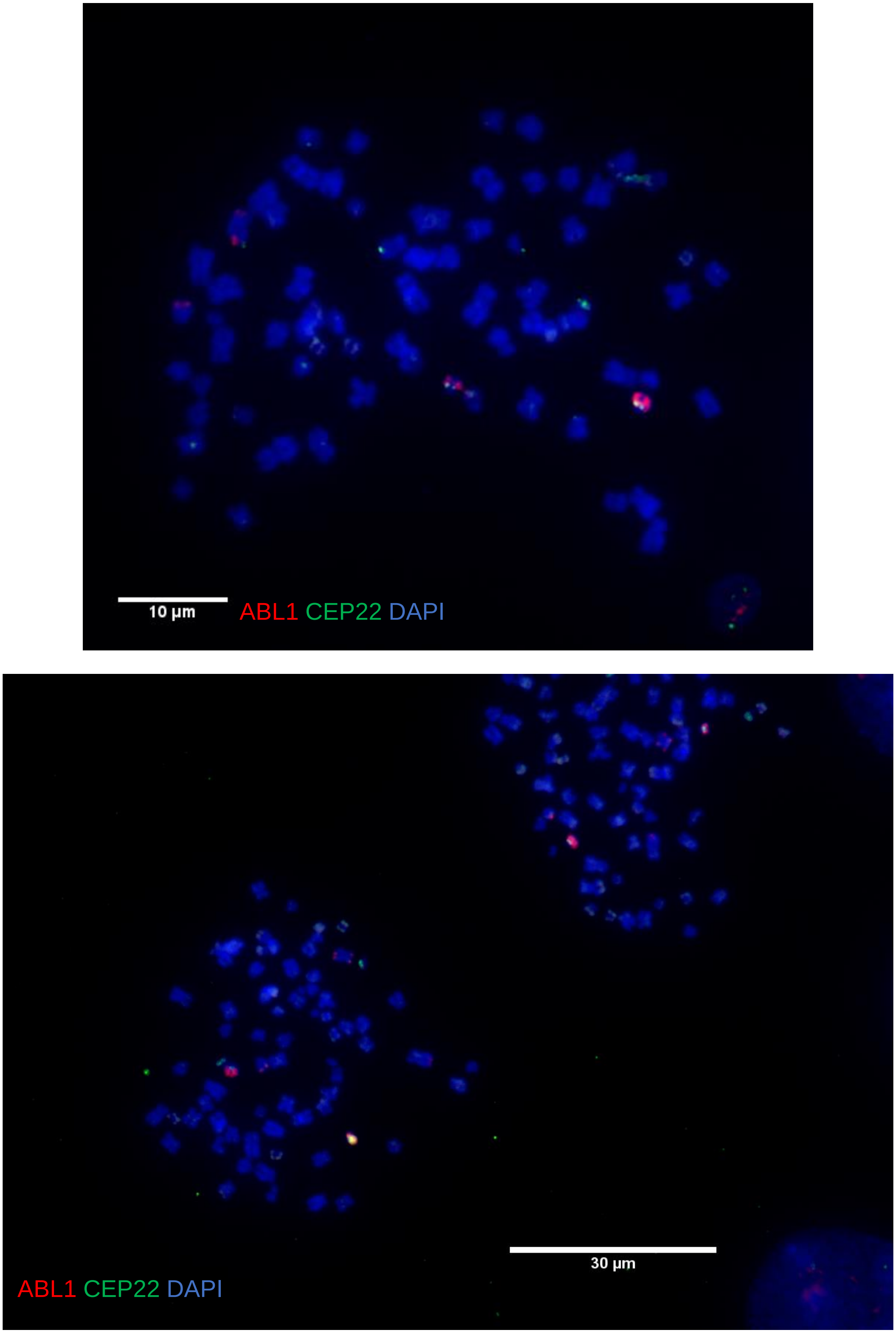


**Figure S7.**

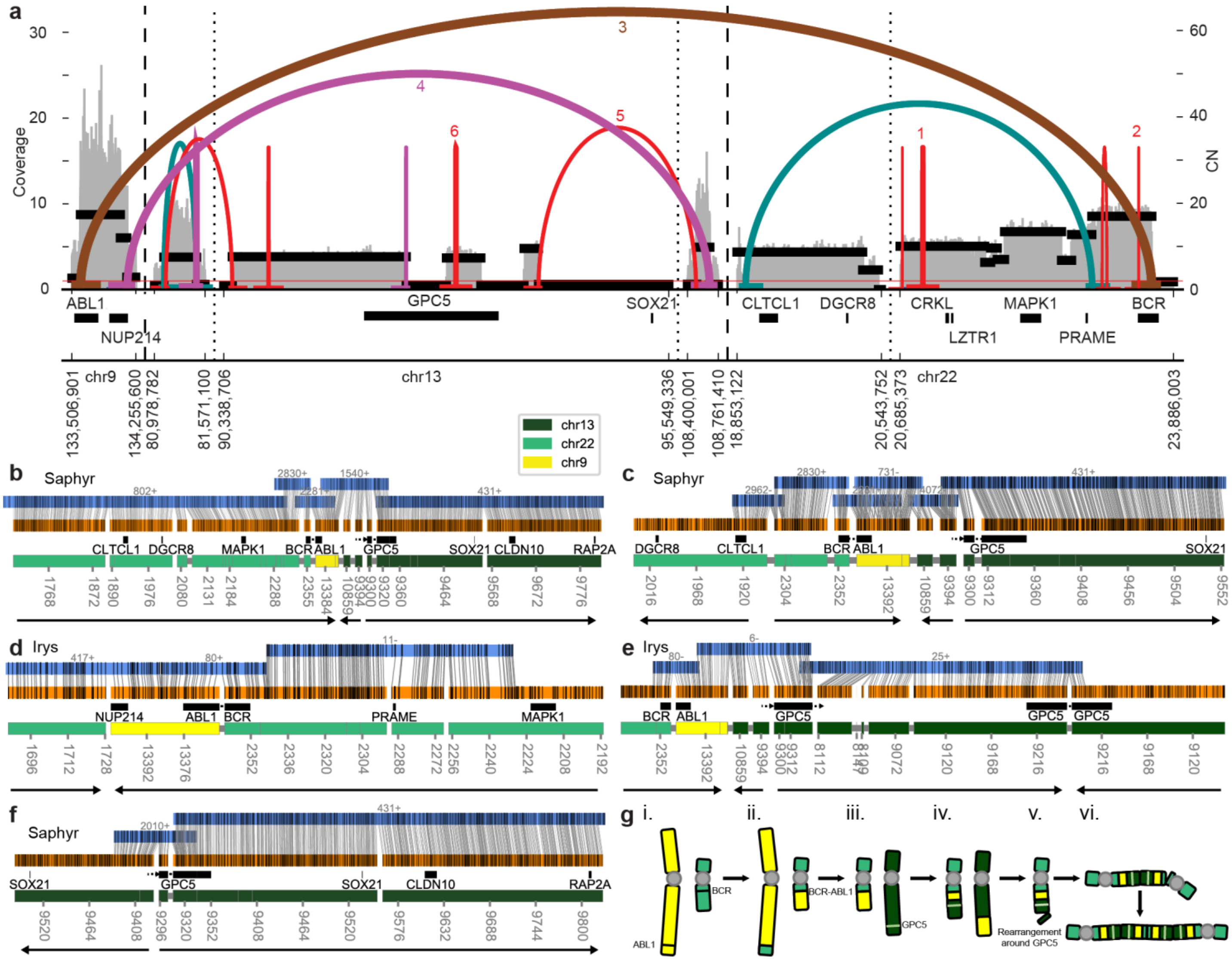
a
Coverage
CN
ABL1
NUP214
GPC5
SOX21
CLTCL1
DGCR8
CRKL
LZTR1
MAPK1
PRAME
BCR
chr9
chr13
chr22
chr13
chr22
chr9
b
Saphyr
c
Saphyr
d
Irys
e
Irys
f
Saphyr
g
i.
ii.
iii.
iv.
v.
vi.
BCR-ABL1
Rearrangement around GPC5

**Figure S8.**

**a**

| Segment | A | B | C | D | E |
|---|---|---|---|---|---|
| Size (kbp) | 1024 | 269 | 217 | 752 | 1176 |
| Raw copy number | 20.0 | 41.4 | 64.3 | 53.0 | 9.6 |
| BFB normalized copy count | 2 | 4 | 6 | 5 | 1 |

**b**

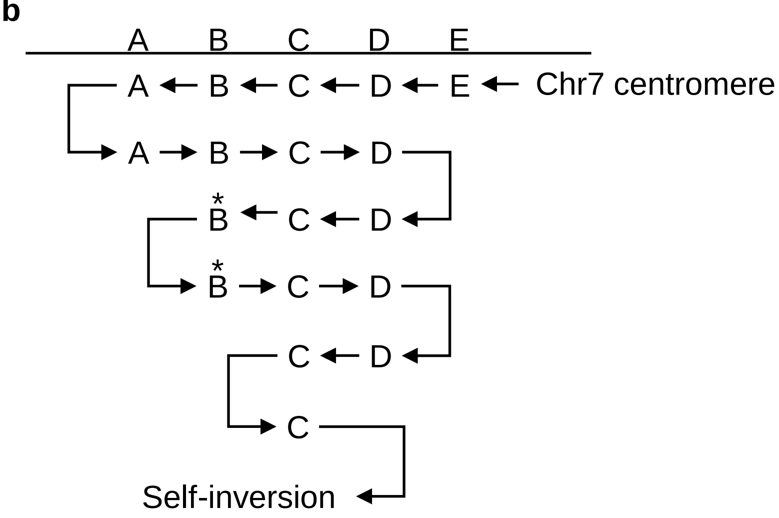


**c**

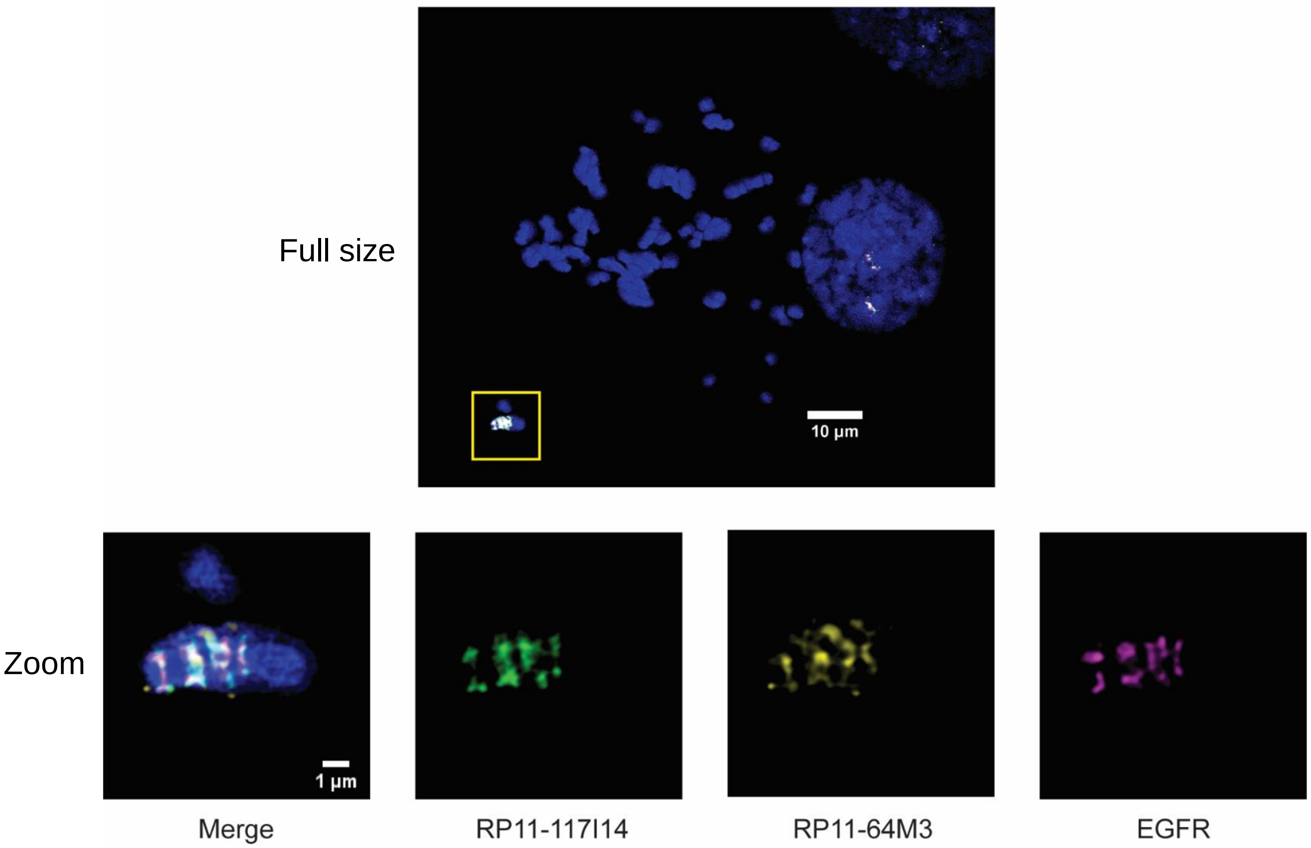


**Figure S9.**

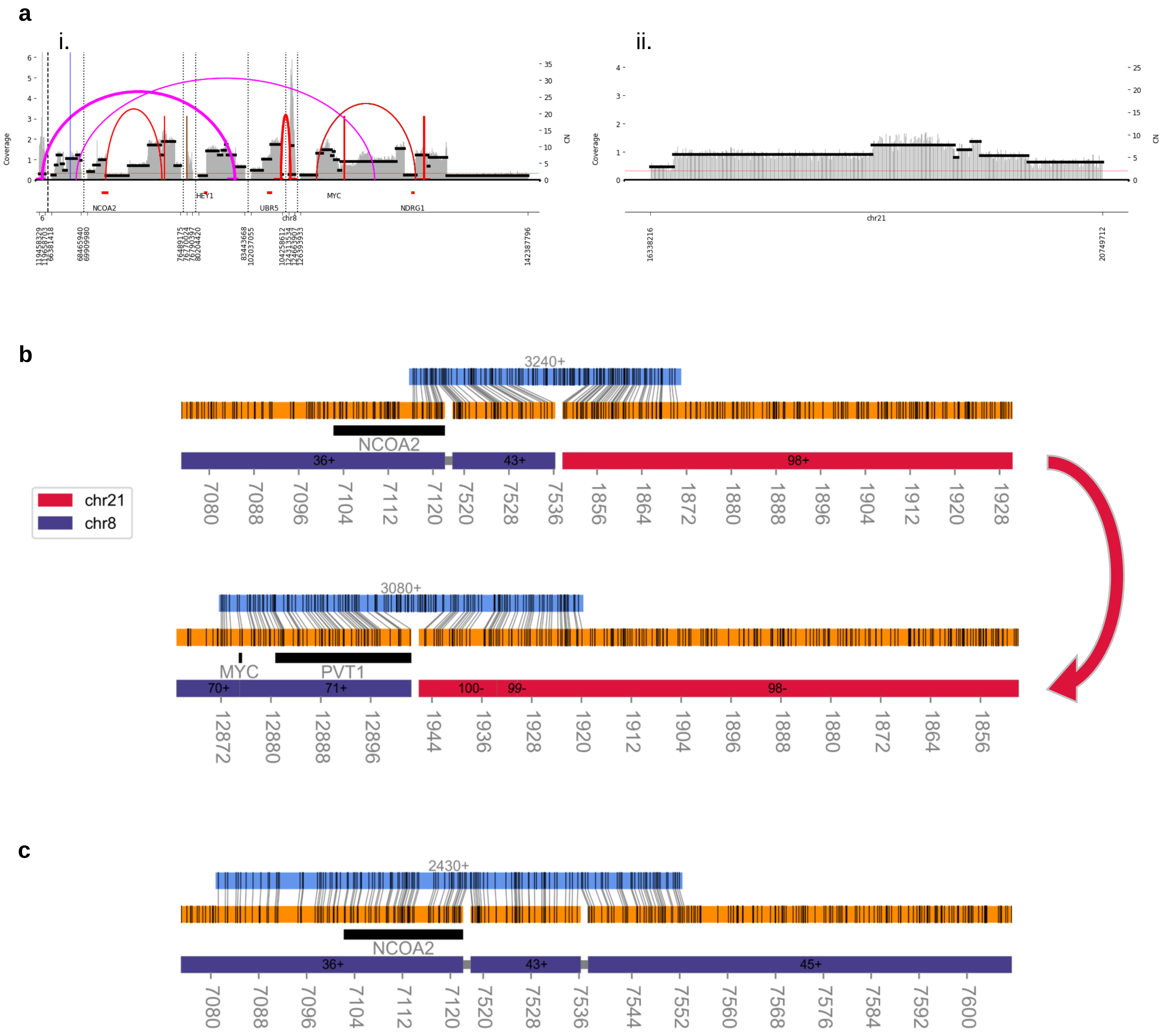
a
i.
ii.
Coverage
CN
NCOA2
HEY1
UBR5
MYC
NDRG1
chr8
chr21
16338216
20749712
142387796
b
3240+
NCOA2
36+
43+
98+
chr21
chr8
3080+
MYC
PVT1
70+
71+
100-
99-
98-
c
2430+
NCOA2
36+
43+
45+

Figure S10.

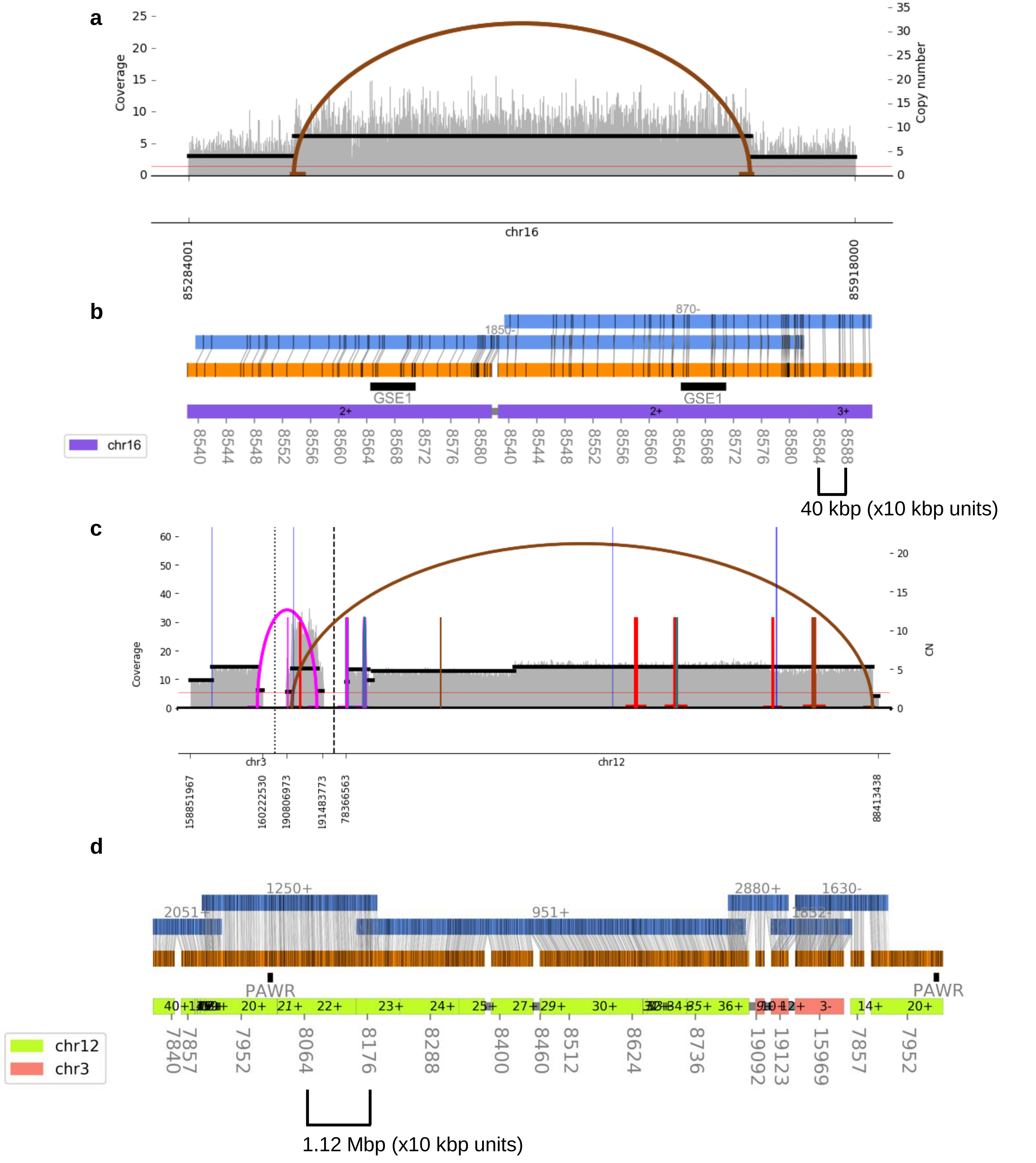
a
Coverage
Copy number
chr16
85284001
85918000
b
870-
1850-
GSE1
GSE1
2+
2+
3+
chr16
40 kbp (x10 kbp units)
c
Coverage
CN
chr3
chr12
158851967
160222530
190806973
191483773
78366563
88413438
d
1250+
2051+
951+
2880+
1630-
1852-
PAWR
PAWR
chr12
chr3
1.12 Mbp (x10 kbp units)

**Figure S11.**
