## Supplemental Figure Captions for "AmpliconReconstructor: Integrated analysis of NGS and optical mapping resolves the complex structures of focal amplifications in cancer"

**Supplemental Figure S1**:

**a**, Long-range sequencing enables accurate disambiguation of large duplications. A breakpoint graph is represented on the left, with colored arrows representing oriented genomic segments, and with edges connecting the segments between source (+) and destination (-) breakpoints. The reconstructions on the right demonstrate that a graph containing duplicated segments can produce multiple possible reconstructions. Short reads provide information about individual junctions in the graph, but long-range sequence information can span and disambiguate multiple junctions. **b**, Diagram of optical map alignment between an OM contig and a reference segment. Unmatched labels prior to the current iteration's best preceding pairings are indicated in red. The SegAligner dynamic programming (DP) recurrence is shown for the matching regions [i,j] and [p,q]. **c**, The SegAligner method for computing statistically significant scoring alignments by estimating parameters in an E-value model from alignments of segments against all contigs present in a sample. An all vs. all representation of best alignment scores of genomic reference segments and OM contigs is shown on the left. The collection of scores of each reference segment are used to build a scoring distribution for the E-value model. In the diagram on the right, alignments are generated from the segment-contig pairings with significantly high-scoring alignment scores. **d**, Ratio of matching regions (MRs) to false-negative reference labels in BioNano NA12878 data indicates the distance-dependent rate of label collapse for consecutive labels in the hg19 reference genome. Two distinct patterns of label collapse exist between BioNano Irys (left) and BioNano Saphyr (right) platforms. Red boxes indicate the regions in the respective technologies that are considered by SegAligner to have non-zero probability of label collapse.

**Supplemental Figure S2:**

**a**, Diagram of features in CycleViz visualization and associated feature tracks. **b**, CycleViz can produce both cyclic (top) and linear (bottom) visualizations. **c**, ScaffoldGraphViewer visualization of AR scaffold graphs using CytoscapeJS. Grey edges indicate connections in the scaffold directed acyclic graph (DAG). Dotted edges indicate imputation. Blue edges indicate edges used in the scaffold’s heaviest path. Green edges indicate a link between two scaffolds.

**Supplemental Figure S3:**

Cumulative precision and recall curves measuring performance of AR reconstructions on simulated amplicon OM data using different OM alignment methods and precision/recall measurement methods. OM assembly failures were not filtered from this figure.

**Supplemental Figure S4:**

**a,** Cumulative precision and recall curves measuring the performance of AR, using SegAligner for OM alignment, on simulated OM data with added false positive edges into simulated amplicon breakpoint graphs. OM assembly failures were not filtered from this panel. **b,** Cumulative precision and recall curves measuring the performance of AR using SegAligner for simulated heterogeneous mixtures of similar amplicons. OM assembly failures were not filtered from this panel.

**Supplemental Figure S5:**

**a**, AA detects a 1.29 Mbp amplified region of chr7 bearing EGFR, and indicates three breakpoint edges. Both EGFR and EGFRvIII appear to be amplified. **b**, AR (with SegAligner) reconstructs a circular 1.26 Mbp ecDNA in GBM39 using a single BioNano Irys contig. A legend for reading CycleViz AR figures can be found in Supplemental Figure S2a,b. Coordinate units on the labeled figure are scaled by 10 kbp. AR reconstructs a form of EGFR carrying the ~20 kbp vIII deletion, which is supported by the breakpoint graph.

**Supplemental Figure S6:**

**a**, AR reconstruction of BCR-ABL1 amplicon in K562 using BioNano Saphyr data. **b,** AR reconstruction of BCR-ABL1 amplicon in K562 using BioNano Irys data. Amplicon was reconstructed in the reverse direction from panel **a** and includes a graph segment (19) which was not included in the Saphyr reconstruction. **c,** Additional AR reconstruction (using BioNano Saphyr data) from AA-detected amplified segments outside the reconstructions in panels **a** and **b**. **d**, Additional AR reconstruction (using BioNano Irys data) from AA-detected amplified segments outside the reconstructions in **a** and **b**, with the exception of the leftmost segment, 20, which appears at the leftmost end of the Irys reconstruction in panel **b.**

**Supplemental Figure S7**:

FISH confocal microscopy on DAPI-stained metaphase chromosomes in K562 cells. Centromeric repeat probe for chr22 (CEP22) and ABL1 are stained in green and red, respectively. Two separate plate regions are shown.

**Supplemental Figure S8:**

**a**, Reproduction of Fig. 3a with SOX21 additionally labeled. **b**, Extended AR reconstruction of K562 BCR-ABL1 amplicon using Saphyr data. **c**, AR reconstruction with Saphyr data supporting an alternate orientation for the genomic segment containing CLTCL as opposed to panel **b**. **d**, AR reconstruction with Irys data containing BCR-ABL1 flanked by chr22 on both sides. **e**, AR reconstruction with Irys data containing BCR-ABL1 with an inverted repeat of the 3' end of GPC5. **f**, AR reconstruction with Saphyr data shows inverted repeat of genomic segment containing SOX21. **g**, Hypothetical model for BCR-ABL1 amplification in K562, involving the reciprocal translocation of q arms of chromosomes 9 and 22 to form the BCR-ABL1 fusion (i-ii), then translocation of q arm of chromosome 13 with the BCR-ABL1 chromosome (iii-iv) with rearrangement of the translocated regions from chr13 on the BCR-ABL1 chromosome (v). Finally, inverted repeats observed in AR scaffolds can be explained by amplifications of BCR-ABL1 through a dicentric chromosome model of amplification (vi).

**Supplemental Figure S9:**

**a**, Table of BFB amplicon segments showing size, raw copy number (as estimated by AA), and normalized BFB count. **b**, A valid BFB string supported by both AA and AR, which contains interior structural variation (marked with an asterisk symbol, "*") **c**, Multi-FISH using super-resolution confocal microscopy on DAPI-stained metaphase chromosomes in HCC827. Brightness was decreased using ImageJ between full size and zoomed images. Probe RP11-64M3 corresponds to segment A, RP11-117I14 corresponds to segment C, and EGFR corresponds to segment D.

**Supplemental Figure S10:**

**a**, Two of five additional AA-generated breakpoint graphs for HCC827. The graph on the left (i) shows a complex structure of rearrangements, while the graph on the right (ii) shows distinct copy number changes without breakpoint graph edges. **b**, AR reconstruction of BioNano Saphyr scaffold joining segments from the graph labeled i in panel **a**, including NCOA2 and MYC, to a segment of chr21 from the graph labeled ii in panel **a.** The scaffold on top in panel **b** can be joined to the scaffold on the bottom as the segment labeled 98 overlaps, as indicated with the curved arrow. **c**, An alternate scaffold reconstructed by AR showing NCOA2 joined to additional segments from chromosome 8.

**Supplemental Figure S11:**

**a**, AA-generated breakpoint graph visualization showing a 430 kbp amplified region with a single breakpoint edge connecting the ends of the amplicon. **b**, AR reconstruction of the amplicon shows a 967 kbp region capturing the segmental tandem duplication involving GSE1. **c**, AA-generated breakpoint graph visualization showing a complex amplified rearrangement joining segments (12.0 Mbp in total) from chr12 and chr3. **d**, AR-reconstructed 13.1 Mbp amplicon containing segments from chr12 and 3, including interior deletions and multiple copies of PAWR.
